# COMPARISON OF UNIVARIATE AND MULTIVARIATE REFERENCE INTERVAL METHODS

**DOI:** 10.1101/2024.11.04.621828

**Authors:** Esra Kutsal Mergen, Sevilay Karahan

**Affiliations:** Hacettepe University Faculty of Medicine, Department of Biostatistics, Ankara, Turkey

**Keywords:** Mahalanobis Distance, Reference Interval, Multivariate Reference Region, Univariate Reference Interval, Simulation Study

## Abstract

**Background:** In clinical practice, reference intervals play a pivotal role in interpreting laboratory test results. Yet, when several tests are taken into consideration simultaneously, the traditional univariate intervals might not suffice due to the elevated risk of Type 1 errors.

**Methods:** This study introduces and evaluates two multivariate reference interval techniques: one based on Mahalanobis distance and the other an adaptation of the multivariate confidence interval. Using Monte Carlo simulations, we focused our assessments on the interplay between “Serum Ferritin and Transferrin Saturation” values.

**Results:** Upon evaluation, it became evident that the multivariate methods significantly reduced false positives. They presented enhanced accuracy over traditional univariate intervals. Notably, the method involving Mahalanobis distance stood out in terms of efficacy.

**Contributions:** Beyond presenting novel techniques, our research underscores the importance and potential of using multivariate approaches in clinical lab settings. The findings can guide better medical decision-making, ensuring optimized allocation of healthcare resources.

**Highlights:**

- We present two alternative multivariate reference interval methods that consider the relationship between two analytes simultaneously.
- Through comprehensive simulation studies, we compare the proposed methods with the conventional univariate reference interval method.
- Our results show the superior performance of our proposed methods in terms of patient rate, sensitivity, specificity, and overall accuracy.
- We discuss the practical relevance of these new methods.

## 1. Introduction

Reference intervals are essential tools that healthcare professionals use to interpret laboratory test results to make medical decisions [1-4]. Using reference intervals, physicians evaluate individuals’ laboratory findings to assess disease status and determine the appropriate treatment. For example, if a patient’s ferritin concentration is not within the reference interval, the value is flagged, and the patient may receive further examination or treatment.

How to properly characterize reference intervals has been described by the International Federation of Clinical Chemistry and Laboratory Medicine (IFCC) and the National Committee for Clinical Laboratory Standards (NCCLS) [2]. Reference interval characterization starts with selecting individuals to construct reference intervals, called as reference individuals. Reference individuals should be healthy individuals who are similar to the target patients in other aspects except the disease status. Then measure of the phenotype of interest is obtained from each reference individual. Using statistical analysis, the lower and upper reference limits for this phenotype are obtained over obtained measures. These lower and upper limits constitute the reference interval for the phenotype of interest [5].

As these intervals are directly used in interpreting laboratory results, the quality of the reference intervals can play a significant role in clinical outcomes [6]. The standard application of reference intervals suffers from using reference intervals where each test result is assessed independently. Such intervals are called univariate reference intervals. However, the physician’s decision-making process includes more than one comparison in which different test results related to the clinical situation are considered together [7, 8]. For example, total cholesterol, LDL, and HDL values for the diagnosis of heart disease [9], TSH, and ft4 values for the diagnosis of goiter disease [10], and Ferritin and Transferrin Saturation percentage values for the diagnosis of anemia can be evaluated together [11]. However, the common practice of examining more than one univariate reference interval together often ignores the information that can be obtained from multivariate relationships.

Hence it decreases the accuracy of evaluation by causing increased Type 1 error rates. Therefore, the practice methods’ reliability is subject to questioning [12, 13].

For example, when a single variable is evaluated with a univariate reference interval, the Type 1 error rate *α* is 0.05. In other words, a healthy individual has (1 − *α*)^1^ probability of being classified as healthy, that is, a 95% probability. However, when the same healthy individual is evaluated with the univariate reference intervals calculated separately for two interrelated variables, she will be classified as healthy with (1 − α)^2^ probability, that is 90.25%. This indicates that the probability of false positive observations increases with the number of univariate reference intervals evaluated [14].

In clinical practice, decreasing the number of false-positive findings, or Type 1 errors, is imperative. Physicians rely on accurate laboratory results to guide further examinations and determine appropriate treatment plans for patients. False-positive results can mislead practitioners into prescribing unnecessary and potentially harmful interventions, thereby exposing patients to undue stress and risk. Moreover, reducing Type 1 errors enables more efficient use of healthcare resources and staff time. Therefore, the reduction in error rates not only enhances patient care but also contributes to lowering overall healthcare costs [6, 15].

Use of multivariate reference intervals is suggested to address false-positive findings that are due to the univariate reference intervals. A seminal study on this topic suggests that when dealing with two related analytes, an ellipse covering the bivariate reference interval could represent a multivariate reference interval [13]. However, this method has not been implemented in clinical biochemistry and laboratory applications due to difficulties in obtaining and interpreting reference limits corresponding to the ellipse in the bivariate reference interval model [16, 17].

In this study, we present two alternative multivariate reference interval methods that consider the relationship between two analytes simultaneously. One of the methods that we propose is based on the Mahalanobis distance and the other is based on the adaptation of the multivariate confidence interval formulation to the multivariate reference interval setting. Through simulation studies, we show that the proposed methods reduce the probability of false-positive findings and overcome challenges in interpreting reference limits corresponding to the elliptical region suggested in 13.

## 2. Methods

### 2.1. Univariate Reference Intervals

Univariate Reference Intervals assume that the distribution of reference values follows a normal distribution. Under the assumption of normality, the univariate reference interval limits are given by equations 1 and 2.

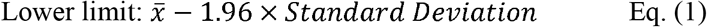

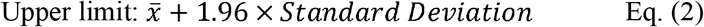

In the equation, 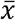 represents the mean of the reference values derived from the reference sample group, while ‘standard deviation’ denotes the standard deviation of the same group. It’s important to note that these calculations operate under the premise that 95% of the reference sample group is healthy [18, 19]. The factor 1.96 is derived from the standard normal distribution table, corresponding to a 95% confidence level.

### 2.2. Multivariate Reference Intervals using Mahalanobis Distance

When constructing multivariate reference intervals, it’s crucial to accurately capture the interrelationships between correlated tests. The Mahalanobis distance stands out as it inherently considers these correlations, unlike standard Euclidean distance. By utilizing the covariance structure of the data, it offers a more genuine representation of multivariate relationships. Moreover, it’s scale-invariant, making it apt for comparing tests with different units. Overall, for correlated clinical tests, Mahalanobis distance ensures a more accurate and clinically relevant assessment of multivariate distances.

Mahalanobis distance measures the distance of an observation from the data’s mean vector in a multidimensional space. This method considers the inter-variable relationships through the variance-covariance matrix, yielding distinct distance scores for each observation [20]. The Mahalanobis distance method is based on the assumption that the data is multivariate normally distributed. The multivariate normal distribution is a generalization of the univariate normal distribution for the number of variables p≥ 2 [21, 22]. Furthermore, this distance follows a pattern known as the chi-square distribution. To determine its significance, the calculated Mahalanobis distance for each data point is then compared with a standard chi-square table value [23].

In this paper, we consider two interrelated tests, hence p=2. Within this context, Σ represents a covariance matrix with dimensions, while x stands for a vector of dimension. Similarly, μ is the mean vector with the same 2×1 dimensions. The formula to compute Mahalanobis distance is given by Eq 3.

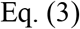

For example, in Figure 1, the observation “A” is inside the ellipse. Therefore, the Mahalanobis distance from “A” to the center is small. Accordingly, “A” is not identified as a multivariate outlier. However, “A” is a univariate outlier for both coordinates. The observation “B” is outside the ellipse. Therefore, the Mahalanobis distance from “B” to the center is relatively large. Thus, the observation is classified as a multivariate outlier. In this method, since the number of variables of interest is two, the chi-square table value is 5.991. Therefore if the calculated Mahalanobis distances of the observations are greater than 5.991, the observation is classified as a patient; Less than 5,991 is classified as healthy.

**Fig 1.**
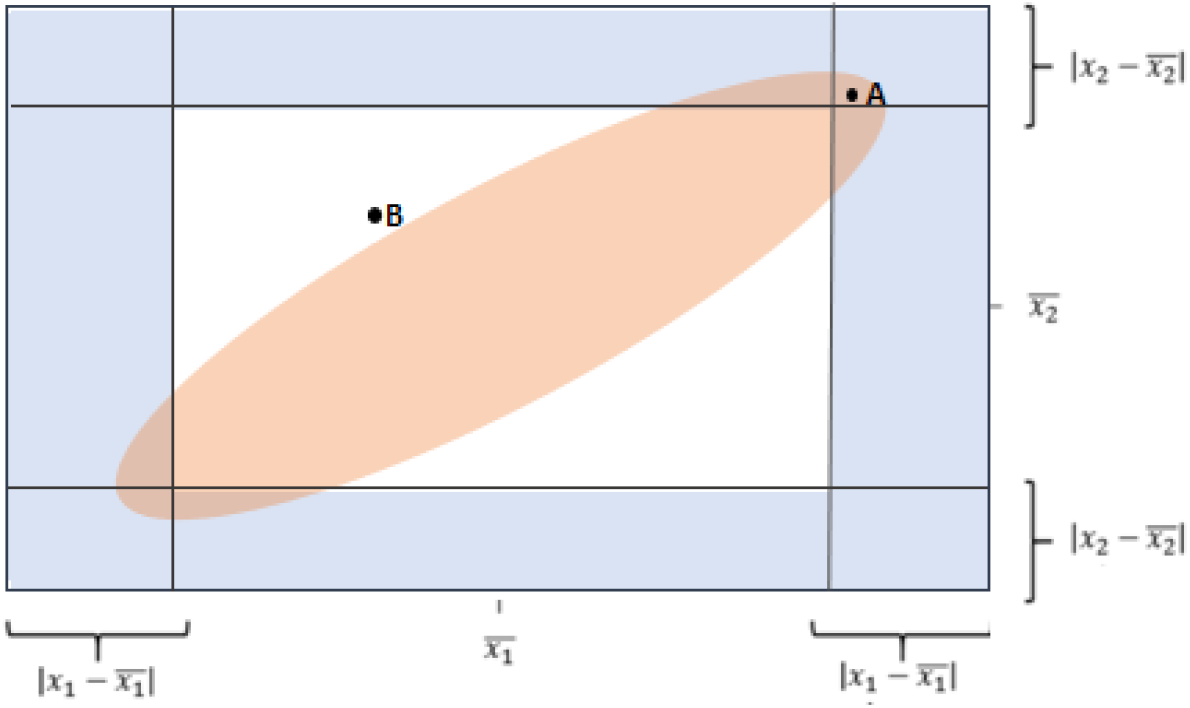
A visualization to illustrate the Mahalanobis method

**Fig 2.**
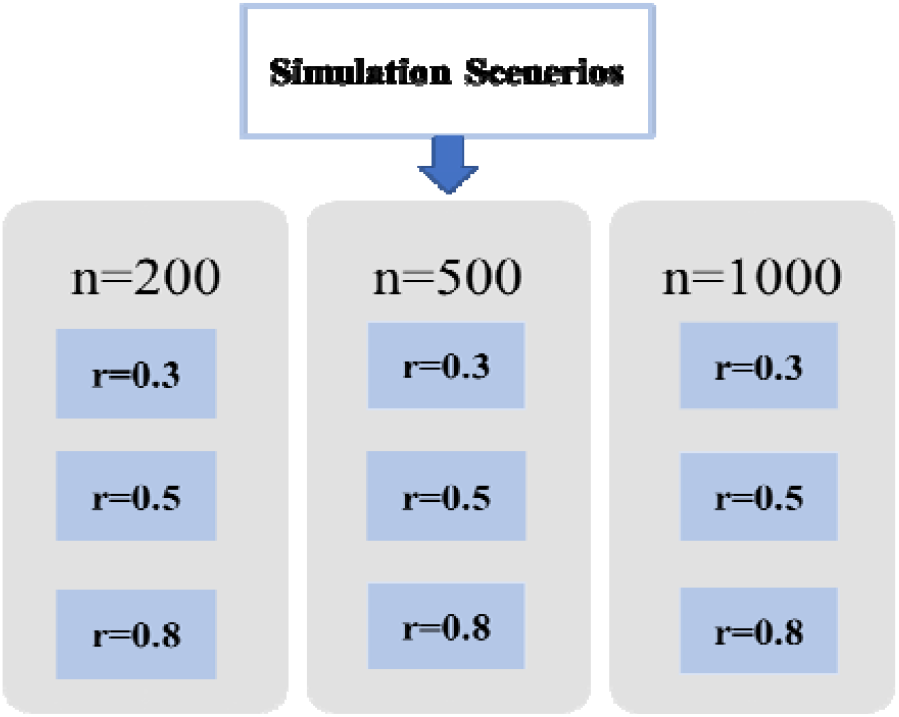
Simulation Scenarios

### 2.3. Multivariate Reference Intervals using Multivariate Confidence Intervals

In this approach we adapt multivariate confidence intervals to determine reference intervals. The confidence interval itself captures variability. It provides a range of values within which the true parameter is expected to lie with a certain level of confidence. In a multivariate context, In a multivariate context, multivariate confidence intervals (MCIs) capture the variability of each parameter separately.

In this setting, we end up with two distinct reference intervals because we’re dealing with two specific variables. To determine the multivariate reference interval limits, we use the mean value of the observations, their standard deviation, and the appropriate values from the chi-square table, all tailored to the number of variables (p) being considered.

The multivariate confidence interval was adapted to the multivariate reference interval formulation. The lower and upper bounds are calculated for two variables as follows:

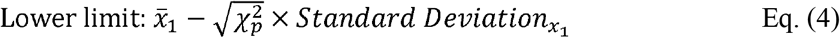

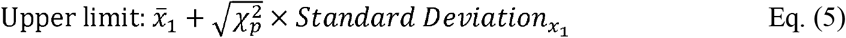

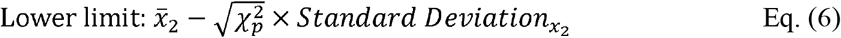

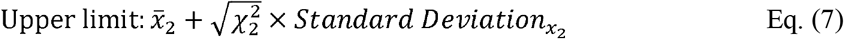

In the method of adapting the multivariate confidence interval to the reference interval formulation, two separate reference intervals are obtained due to the calculation of two variables. Since Mahalanobis distance considers the relationship between variables and is chi-square distributed, the relevant table value was used for the number of p variables. Multivariate reference interval limits are obtained using the observations’ mean, standard deviation, and chi-square table values based on variables.

## 3 Simulation and Results

In this section, we present a simulation study to evaluate and compare our proposed multivariate reference interval methods—based on the Mahalanobis distance and multivariate confidence intervals—against the conventional univariate reference interval method. Using the R software, we examined various scenarios, considering different correlation coefficients (r) and sample sizes (n). Our central focus in this comparison is on four metrics: the rate of patient classification, sensitivity, specificity, and overall accuracy. Through this study, we aim to highlight the potential benefits and superior performance of the new techniques when contrasted with the conventional univariate reference intervals.

We created a synthetic dataset using Monte Carlo simulation to mirror real-world lab test results. Our focus was on “Serum Ferritin and Transferrin Saturation” values, which are vital in diagnosing anemia. We based our dataset’s statistics, specifically the mean and standard deviation for the two values, on the findings from the research article “Reference values for serum ferritin and percentage of transferrin saturation in Korean children and adolescents” [24]. It’s important to note that in our dataset, males and females aged 10-20 have separate mean and variance values, reflecting the differences seen in the real world. Finally, our data was generated following a bivariate normal distribution, ensuring it’s representative of genuine lab data structures.

In this study, we introduced specific terminologies to represent different methods of categorizing instances based on their measurements. Specifically, the term “Uni1” denotes a method where instances are labeled as patients if measurements for either of the studied variables fall outside the univariate reference intervals introduced in Section 2.1., following the “or” rule. Conversely, “Uni2” refers to a method where instances are categorized as patients only when measurements for both studied variables exceed the univariate reference intervals, applying the “and” rule.

We utilize a method termed ‘Mahalanobis’ which is rooted in the Multivariate Reference Intervals that employ the Mahalanobis Distance introduced in Section 2.2. In the context of multivariate reference intervals derived from Multivariate Confidence Intervals introduced in Section 2.3., “Multi1” is a method that identifies instances as patients if their measurements deviate from the multivariate intervals for either of the variables, according to the “or” rule. On the other hand, “Multi2” recognizes instances as patients when their measurements surpass the multivariate range for both variables, employing the “and” rule.

We conducted simulations for various scenarios, considering different sample sizes (n = {200, 500, 1000}) and correlation coefficients (r = {0.3, 0.5, 0.8}) as depicted in Figure Datasets following a bivariate normal distribution were generated for each simulation scenario. Classification was performed using the above methods (Uni1, Uni2, Multi1, Multi2, Mahalanobis). For each scenario, we conducted 1000 replications and calculated the patient rates as well as sensitivity, specificity, and overall accuracy. Note that here the Mahalanobis method in Section 2.2. was accepted as the gold standard.

Table 1 shows the patient rates of the methods under different scenarios for males and females aged 10-20. In general, as the correlation between the two variables of interest increases, the rate of patient classification decreases in the Uni1 method for both subgroups, while the rate increases in the Uni2 method. On the other hand, when the sample size increases while the correlation coefficient is constant, the patient classification rates do not show a significant difference in the Uni1 and Uni2 methods. The proposed Mahalanobis method provided an approximately 5% patient classification rate for both subgroups in all scenarios with different sample sizes and different correlation structures. In the Multi1 method, the rate of patient classification decreases as the correlation between the variables increases for both subgroups. In the Multi2 method, as the correlation between the variables increases, the rate of patient classification increases for both subgroups.

**Table 1.** Patient rates of the methods for males and females aged 10-20 under different scenarios.

| Gender | Correlation | Method | Diseased Rate (%) |  |  |
| --- | --- | --- | --- | --- | --- |
|  |  |  | n=200 | n=500 | n=1000 |
| Female | 0.3 | Uni1 | 9.33 | 9.44 | 9.48 |
|  |  | Uni2 | 0.46 | 0.49 | 0.48 |
|  |  | Mahalanobis | 4.95 | 4.99 | 4.98 |
|  |  | Multi1 | 2.77 | 2.80 | 2.78 |
|  |  | Multi2 | 0.06 | 0.07 | 0.07 |
|  | 0.5 | Uni1 | 8.94 | 9.02 | 9.03 |
|  |  | Uni2 | 0.88 | 0.92 | 0.90 |
|  |  | Mahalanobis | 4.97 | 4.99 | 4.97 |
|  |  | Multi1 | 2.68 | 2.70 | 2.69 |
|  |  | Multi2 | 0.17 | 0.16 | 0.15 |
|  | 0.8 | Uni1 | 7.71 | 7.74 | 7.80 |
|  |  | Uni2 | 2.17 | 2.17 | 2.20 |
|  |  | Mahalanobis | 4.97 | 4.99 | 5.00 |
|  |  | Multi1 | 2.37 | 2.35 | 2.30 |
|  |  | Multi2 | 0.52 | 0.51 | 0.50 |
| Male | 0.3 | Uni1 | 9.36 | 9.49 | 9.48 |
|  |  | Uni2 | 0.46 | 0.48 | 0.49 |
|  |  | Mahalanobis | 4.91 | 4.99 | 4.99 |
|  |  | Multi1 | 2.74 | 2.79 | 2.81 |
|  |  | Multi2 | 0.07 | 0.07 | 0.07 |
|  | 0.5 | Uni1 | 9.06 | 9.03 | 9.08 |
|  |  | Uni2 | 0.90 | 0.91 | 0.92 |
|  |  | Mahalanobis | 4.97 | 4.99 | 4.97 |
|  |  | Multi1 | 2.71 | 2.70 | 2.73 |
|  |  | Multi2 | 0.19 | 0.17 | 0.19 |
|  | 0.8 | Uni1 | 7.73 | 7.74 | 7.75 |
|  |  | Uni2 | 2.18 | 2.20 | 2.21 |
|  |  | Mahalanobis | 4.97 | 4.99 | 4.98 |
|  |  | Multi1 | 2.32 | 2.34 | 2.32 |
|  |  | Multi2 | 0.52 | 0.51 | 0.53 |

In all methods, as the sample size increases, the variability of the number of patients decreases as expected. In all scenarios, the average percentage of patient classification was found highest in the Uni1 method and the lowest in the Multi2 method. In contrast, the Multi1 method had approximately a similar patient ratio to the Mahalanobis method.

The patient rates obtained from our simulations are visually represented in Figure 3 and 4 using box plots. Upon examining these graphs, several trends become evident. In the Uni1 method, as the correlation coefficient between the variables increases, the patient rate value tends to decrease. In contrast, the Uni2 and Multi2 methods show an increase in patient rates and greater variability as the correlation coefficient increases.

**Fig 3.**
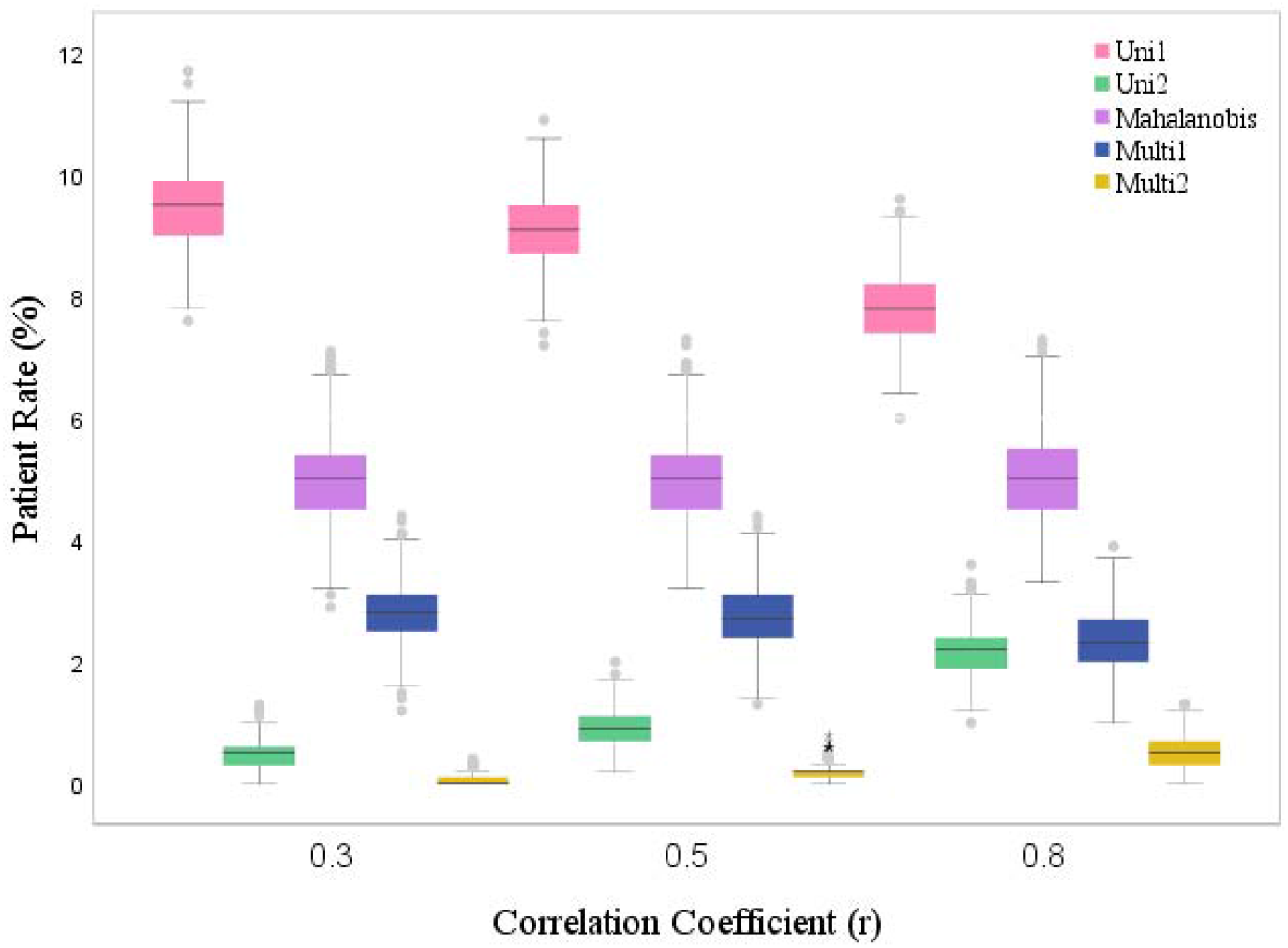
Comparison of the patient rates of the methods in case the sample size is 1000 and the subgroup is female under different scenarios.

**Fig 4.**
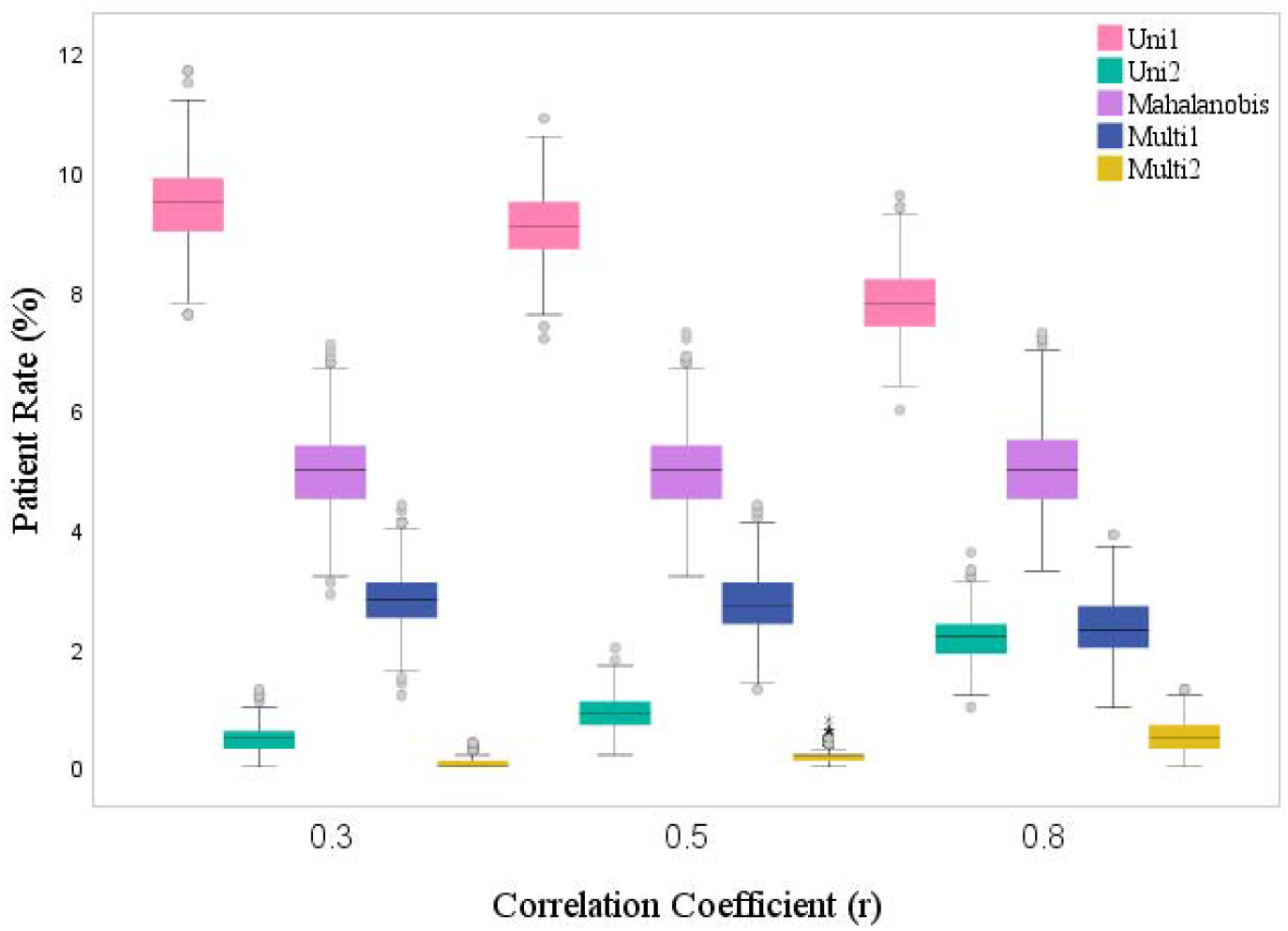
Comparison of the patient rates of the methods when the sample size is 1000 and the subgroup is male under different scenarios.

For the Multi1 method, there is a slight decrease in the patient rate as the correlation value increases. However, in the Mahalanobis method, the patient rate remains relatively stable regardless of changes in the correlation coefficient. Furthermore, the patient rate variability decreases in the Mahalanobis method as the correlation coefficient increases, indicating a more consistent performance across different scenarios.

In our analysis, we designated the Mahalanobis method as the benchmark, against which we measured the performance of the other methods, calculating their sensitivity, specificity, and overall accuracy. The results, presented in Table 2, reveal that the Multi1 method consistently exhibits the highest accuracy and specificity across all scenarios. Moreover, the sensitivity value of the Multi1 method closely aligns with that of the Uni1 method, although it’s worth noting that the sensitivity and specificity values can be influenced by the underlying prevalence of the condition [25].

**Table 2.** Sensitivity, specificity, and accuracy for different correlation coefficients when the Mahalanobis method is accepted as the benchmark (n=1000)

| Correlation | Method | Sensitivity (%) | Specificity (%) | Accuracy (%) |
| --- | --- | --- | --- | --- |
| 0.3 | Uni1 | 87.16 | 94.76 | 94.39 |
|  | Uni2 | 8.15 | 99.97 | 95.46 |
|  | Multi1 | 55.97 | 100 | 97.86 |
|  | Multi2 | 1.13 | 100 | 95.14 |
| 0.5 | Uni1 | 81.24 | 95.02 | 94.32 |
|  | Uni2 | 14.68 | 99.84 | 95.55 |
|  | Multi1 | 53.89 | 100 | 97.67 |
|  | Multi2 | 3.25 | 100 | 95.12 |
| 0.8 | Uni1 | 68.09 | 95.56 | 94.21 |
|  | Uni2 | 27.11 | 99.19 | 95.64 |
|  | Multi1 | 47.48 | 100 | 97.41 |
|  | Multi2 | 9.49 | 100 | 95.55 |

Since reference intervals are statistically derived tools used to characterize the central 95% of the healthy population, the approximately 5% patient ratio achieved in each scenario with the Mahalanobis method proves that this method can be recommended in the face of the uncertainty of the multivariate reference region and the difficulty of implementation. Furthermore, although Mahalanobis gives the closest patient classification rate of 5%, the Multi1 method, which offers the most comparable results to this method, will facilitate clinical evaluation since it does not give the lower and upper reference limits that the physician can quickly evaluate.

## 4. Discussion

In the realm of medical diagnostics, the utilization of reference intervals, particularly univariate reference intervals, has remained a fundamental practice for the interpretation of laboratory results. This study has primarily focused on the development and assessment of methods that are based on multivariate reference intervals using mahalanobis distance and multivariate confidence intervals. Both require comprehensive studies involving measurements obtained from a carefully selected population of “healthy” individuals. It is imperative to ensure that individuals chosen for reference studies are devoid of acute or chronic illnesses, refrain from medications that may influence laboratory test outcomes, and can carry out their routine daily activities [18].

It is a prevailing situation for physicians to have one or more results that are minimally outside the reference interval from the laboratory values routinely requested from clinically healthy individuals. Considering that a healthy individual has a 5% chance of having a result value outside the reference limits, and this chance is valid for every desired laboratory test result, the probability of a healthy individual’s test result being outside the reference interval will increase as the number of tests increases. This will increase the false positive rate. In such a situation, the multivariate reference interval method was proposed as an alternative to univariate reference interval methods in the clinical laboratory literature over thirty years ago [23],[26]. Early studies on this subject summarized the advantages of the multivariate reference region method in reducing the number of “false positive” results found using univariate reference intervals [27, 28]. However, the difficulties in obtaining and interpreting the multivariate reference interval as a region due to increased dimensionality have also been noted in studies [29]. Few studies have been published on the multivariate reference region approach to date. The existing multivariate reference interval approaches cannot be applied in clinical practice because of their dimensionality [30-37]. Specifically, they are not easy to calculate in clinical practice and to be understood and interpreted by physicians. Therefore, it is unsurprising that the existing multivariate reference interval approaches have not been applied in this area, playing only a marginal role, as noted by Harris and Boyd and Wright and Royston [30, 38]. These factors underscore the existence of unexplored methods, hence developing intuitive multivariate reference interval techniques is a promising avenue for research.

In practice, it will be a great advantage to present the multivariate reference interval approach as not a region to help physicians easily apply it. The limitation of this study is that it considers the data distributed according to bivariate normal distribution. For non-normally distributed variables, a transformation can be made by following the approaches outlined by Harris and Boyd [38]. Approaches to nonparametrically construct multivariate reference interval methods have been described in various studies [38, 39]. In future studies, Multivariate reference interval approaches should be developed for situations where there are more than two variables, negative correlation structure between the variables, and the variables are not suitable for the normal distribution structure.

## Acknowledgements

This study was supported by the Hacettepe University BAP Coordination Unit.

## 5. Research funding

This work has been supported by Hacettepe University Scientific Research Projects Coordination Unit under grant number THD-2021-19616.

